# Numbers of mutations within multicellular bodies: why it matters

**DOI:** 10.1101/2022.09.26.509555

**Authors:** Steven A. Frank

**Affiliations:** Department of Ecology and Evolutionary Biology, University of California, Irvine, CA 92697-2525, USA

**Keywords:** Luria–Delbrück, population genetics, genetics of disease, somatic mosaicism, cancer, neurodegeneration

## Abstract

Multicellular organisms often start life as a single cell. Subsequent cell division builds the body. Each mutational event during those developmental cell divisions carries forward to all descendant cells. The overall number of mutant cells in the body follows the Luria–Delbrück process. This article first reviews the basic quantitative principles by which one can understand the likely number of mutant cells and the variation in mutational burden between individuals. A new Fréchet distribution approximation greatly simplifies calculation of likelihoods and intuitive understanding of process. The second part of the article highlights consequences of somatic mutational mosaicism for understanding diseases such as cancer, neurodegeneration, and atherosclerosis.

## Introduction

Multicellular organisms often begin as a single cell. Similarly, microbial populations often expand from a small number of colonizing progenitor cells. As clones expand by cell division, mutations inevitably arise. Mosaic multicellular soma or genetically heterogeneous microbial populations follow.

Somatic mutants affect cancer risk.^1,2^ Mutants in microbial clones initiate the spread of faster growing variants, a tumor-like overgrowth.^3^ In plants, somatic variability may promote defense against pathogens and herbivores.^4,5^

This article provides a simple overview of clonal heterogeneity and its consequences. The first part outlines conceptual and quantitative aspects. The most important idea concerns the high diversity between clones. Most cellular populations have relatively few mutants. Some populations have a lot of mutants.

The second part of this article discuss why it is important to understand the diversity of mutants within and between cellular populations. That part illustrates importance by examples.

The examples include diseases such as cancer, neurodegeneration, and atherosclerosis. Other examples clarify the distinction between how diseases begin and how they spread within bodies. Another key way in which somatic mutation affects disease concerns the strong tendency for variation in risk between individuals.

## The Luria–Delbrück problem

The number of mutants in cellular populations was originally studied to estimate the mutation rate.^6^ By counting mutant cells after exponential growth, one can infer how often new mutations arise. This section introduces quantitative aspects of how mutations accumulate in populations.

Suppose a clone begins with one cell and grows to *N* cells. Ignoring cell death, such growth requires *N* – 1 cell divisions because each division adds one cell to the population. If the mutation rate per cell division is *u*, then the expected number of mutational events is (*N* – 1)*u*, which is approximately *Nu* for large *N*. However, knowing the number of mutational events does not by itself tell us the number of mutant cells.

A single mutational event might happen in the first cell division, causing approximately 1/2 of all cells to carry the mutation, with *N*/2 mutants cells in the final population. Or the mutation might happen in a final cell division, causing only one cell to carry the mutation. We must map each mutational event to its time within the cell lineage hierarchy (Fig. 1).

**Figure 1:**
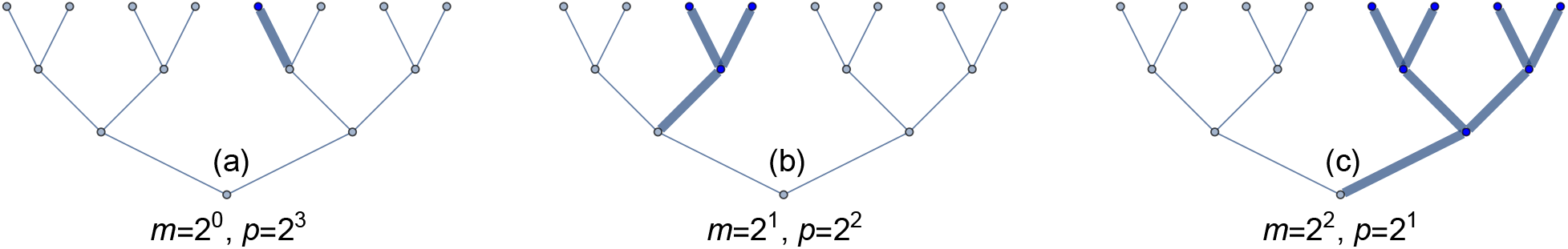
Number of mutants in a growing population depends on the time of the first mutation. (a) When the first mutation happens in the last cell division, then only one cell (*m* = 2°) carries that mutation. In this case, there are *p* = 2^3^ different final cells that could be mutated in the final round of cell division, so such mutations confined to a single cell will be relatively common. (b) When the first mutation happens in the second to last cell division, then two cells (*m* = 2^1^) carry that mutation. In this case, there are *p* = 2^2^ different final groups of cells that could be mutated, so such mutations confined to two cells will be 1/2 as common as for singleton mutations. (c) When the first mutation happens in the third to last cell division, then four cells (*m* = 2^2^) carry that mutation. In this case, there are *p* = 2^1^ different final groups of cells that could be mutated, so such mutations confined to four cells will be 1/4 as common as for singleton mutations. Focusing on single mutational events, every doubling for the final number of mutant cells decreases the probability of occurrence by 1/2.

Deleterious mutations slow the rate of cell division in descendants. Advantageous mutations speed cell division. Such changes alter the final number of mutants. A mathematical literature considers the variety of problems.^7^ We may call the general challenge the Luria-Delbrück problem, named after the initial studies.

I limit the following quantitative sections to a few details about the numbers of neutral mutations. Those calculations provide sufficient introduction to the main conceptual background for interpreting various biological scenarios.

## Average number of mutants

### Intuitive summary

To calculate the average number of mutants in a clone, start by picking one of the *N* final cells. The probability that it carries a mutation is the number of cell divisions that separate that final cell from the initial progenitor cell. The expected number of cell divisions is log(*N*), using the natural logarithm (see next subsection). With a mutation rate of *u* per cell division, the mutant probability per final cell is approximately *u* log(*N*) when the mutation rate is sufficiently small. With *N* final cells, we expect an average of approximately

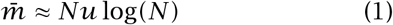

mutants in the population.^7^

A human body has more than *N* = 10^13^ cells.^8^ Estimates for the somatic mutation rate per gene per cell division vary.^9–11^ Here, I use *u* = 10^-6^. The average number of cellular generations is log(10^13^) ≈ 3°. So we expect roughly 3 × 10^8^ mutants for each gene and a frequency of mutants of roughly 3 × 10^-5^. That is a lot of mutant cells but a low frequency of mutations per cell.

A human genome has about 2 × 10^4^ genes and also a lot of noncoding DNA that can influence phenotype. When considering all of the various sites that can be mutated, the overall amount of mutation in the final population of cells is large. And, perhaps more importantly, the mutational variation between multicellular bodies tends to be big, as discussed below.

Microbial clones can also grow into large populations, also carrying substantial numbers of mutants and large amounts of variation between clones.

### Details

The expected number of cellular divisions can be derived by noting that, starting with one cell, a final population of *N* arises by continuous exponential growth such that *N* = *e^rt^*. Thus

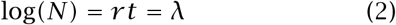

is the expected number of cellular divisions from a final cell back to the original cell, in which *r* is the cellular division rate and *t* is time.

One interpretation is that the actual number of divisions in different cellular lineages follows a Poisson distribution with mean *λ*. Suppose each lineage with *k* divisions contributes to part of a cellular expansion with size 2^*k*^. Summing over the Poisson probabilities *p_k_* for *k* divisions in a lineage is

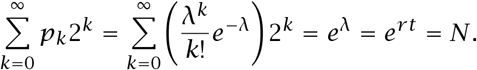

## Variation in number of mutants

The way in which mutations accumulate in growing populations inevitably causes a lot of variation between populations. That large variation may often be more important biologically than the average. For example, disease often concerns what happens in variant or extreme individuals rather than what happens in an average individual. Thinking of the extreme of a probability distribution as what happens in its tail, we may say that the characteristics of a probability distribution’s tail may be the important attributes for a particular problem.

Also, when a distribution has occasional exceptionally large values, the average value does not provide a good sense of what is typical. For example, suppose we observe five sample values, 800, 900, 1000, 1100, 9000. The median is 1000, a good description of central location and typical values, whereas the average is 2560, a poor description of typical values.

## Distributions with fat tails

Figure 1 shows how mutations accumulate in growing populations. When the mutation happens late, as in Fig. 1a, then a small fraction of the population carries that mutation. Such late-occurring mutations happen frequently, because there are more target cell divisions in which a mutation can arise.

For each generation earlier that a mutation might occur, the number of final cells carrying the mutation doubles and the probability of occurrence declines by 1/2, as in Fig. 1b,c. Here, to illustrate, we are simplifying growth to be a sequence of discrete binary cellular divisions.

Consider a large final population of cells, *N*. If a mutation happens in the first cell division of the progenitor cell, then *N*/2 cells are mutated with probability *u,* the probability of mutation during a cell division. When a mutation happens in the second cell division from the original cell, then *N*/4 cells are mutated with probability 2*u*, because there are twice as many target cells. In the third round, *N*/8 cells would be mutated with probability 4*u*. Figure 2a provides an example in which probability doubles with each halving of the number of mutants.

**Figure 2:**
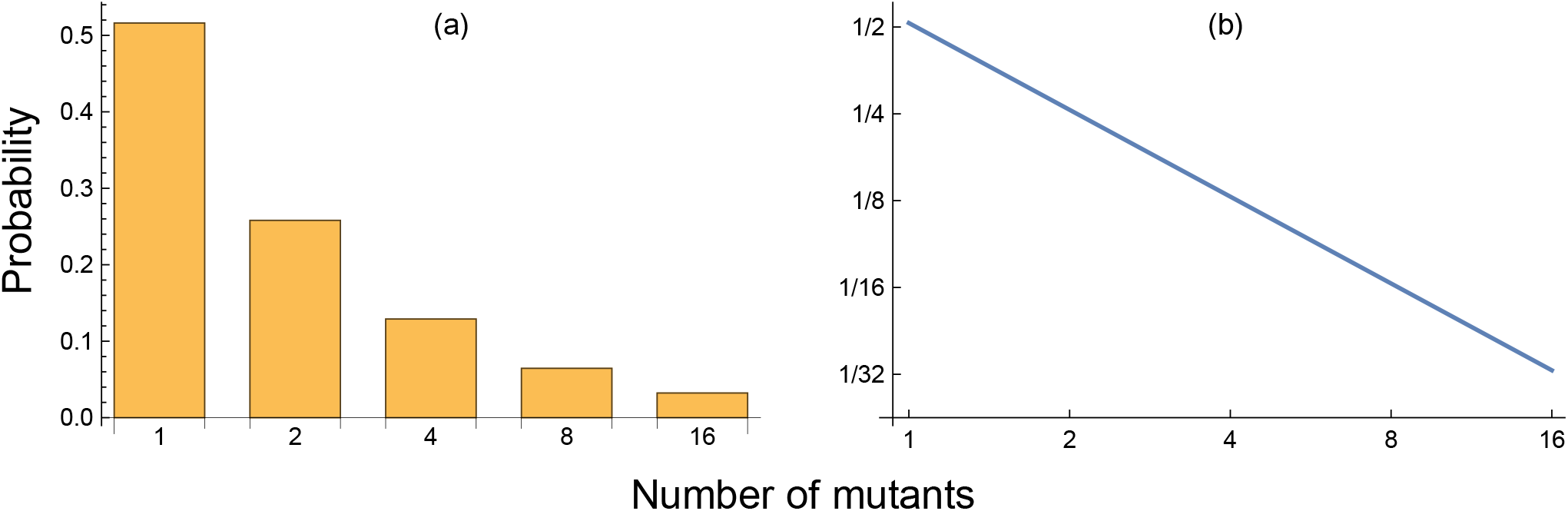
The probability of observing a particular number of mutants in the final population. (a) Probability declines by 1/2 for each doubling in the number of mutants in the final population. (b) A log-log plot of probability versus mutant number has a slope of minus one. This example follows from a discrete binary pattern of cellular division with a single mutational event, as in Fig. 1. The actual log-log slope for a Luria–Delbrück process is approximately - (1 + *e*/2) ≈ −2.36 (from eqn 5). Part of the increase arises because multiple mutational events cause a more rapid decline at the extreme of the earliest mutations and greatest final mutant numbers.

If we make a log-log plot of the probability of occurrence for each number of mutant cells in the final population versus the number of mutant cells, that plot of the upper tail is a straight line with a slope of minus one (Fig. 2b). A straight line of a log-log plot for a probability distribution is a power law. We say that the upper portion of this distribution has a power law tail, or fat tail. The tail is fat because a power law declines relatively slowly, causing a lot of probability and a high chance of observation to occur for large values, in this case, large values for the total number of mutated cells in the final population.

This explanation oversimplifies because it focuses on only a single mutational event. An actual expanding population may have many mutational events. However, mutational events early in the growth of the population are rare because there are few target cells. And those early mutations dominate the upper tail because when they occur they carry forward to all descendant cells. So the simplification does reasonably well for describing the fat upper tail.

This section illustrated the problem by using a discrete binary branching processes for population growth. However, most applications focus on continuously growing populations with overlapping generations, such as in eqn 2. The next section returns to the continuous interpretation.

## Distribution of mutants

No simple formula describes exactly the probability of observing *m* mutants in a final population of size *N*. A large literature provides various ways of making the calculation or of simulating populations with a computer.^7,12^ It is easy to calculate the average, given approximately in eqn 1. But, as noted in the previous section, the distribution has a long fat tail, associated with much variation between populations. It is often the large variation that matters in application rather than the average. I discuss some examples in the second part of this article.

### Fréchet distribution

The prior section noted that the upper tail has a power law shape. But what does the full distribution look like for the typical assumption of continuous population growth? Recently, I found that the distribution of mutants closely matches the well known Fréchet probability distribution.^15^

For a Fréchet distribution, the cumulative probability that the number of mutants is less than or equal to *m* is

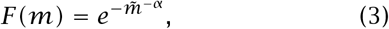

in which 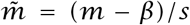. The probability of being in the upper tail, greater than *m*, is 1 - *F*(*m*). The three parameters set the shape, *α*, the scale, *s*, and the minimum value, *β*, such that *m* ≥ *β*.

### Approximate fit

Using the following parameters provides a close match between the Fréchet distribution and the Luria–Delbrück process for neutral mutations

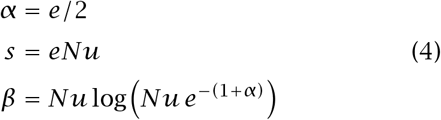

in which *e* is the base of the natural logarithm. The Fréchet match to the Luria–Delbrück depends on the single parameter, *Nu*, the population size multiplied by the mutation rate. Figure 3 shows the match.

**Figure 3:**
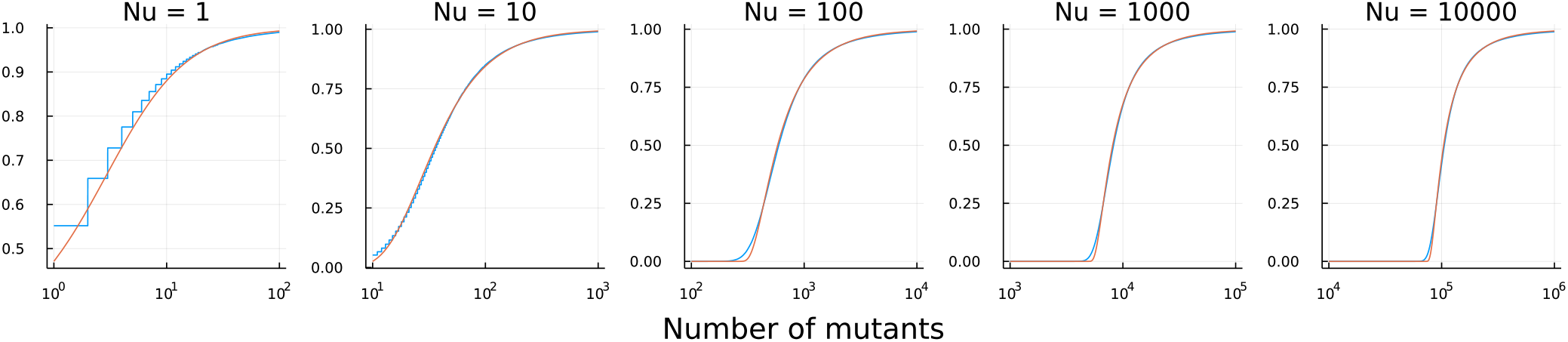
Cumulative probability distribution for the number of neutral mutants. Each population starts with one cell and grows to *N* cells. Mutations occur at rate *u*. The blue curves show the Luria-Delbrück distribution calculated by the simu.cultures computer simulation of the R package rSalvador.^12^ The orange curves show the Fréchet distribution in eqn 3. Reprinted from from Figure 1 of Frank.^15^ See that article for further details. The mathematical reason for the close match between the Luria-Delbrück distribution and the Fréchet distribution arises by linking two separate studies. Kessler & Levine^13^ showed that the Luria-Delbrück distribution converges to a Lévy *α*-stable distribution for large *Nu*. Separately, Simon^14^ showed the close match between the Lévy *α*-stable distribution and the Fréchet distribution. Using the Fréchet distribution provides a benefit because no explicit mathematical expression exists for the Lévy *α*-stable probability distribution.

### Intuition

Why would a Fréchet distribution provide a good match for the number of mutants in a Luria– Delbrück process? To begin, think about the final cells in the population after exponential growth. Look back from those final cells through the multiple rounds of cell division that trace ancestry toward the original progenitor. The mutation farthest back in time from the final cells toward the progenitor typically dominates in determining the number of mutants in the final population.

That most extreme time tends to follow the extreme value probability distribution known as Gumbel.^16–18^ Then, because the most extreme mutation will subsequently expand by multiplicative growth to determine the final number of mutations, we can use the fact that a multiplicative process substituted into a Gumbel distribution leads to a Fréchet distribution.^18^

## Mutation rate

Estimating the mutation rate provided the original motivation to Luria & Delbrück.^6^ Over time people have discovered links between how mutations accumulate in growing populations and topics such as cancer and neurodegeneration. I take up those topics in later sections. This section begins with the mutation rate.

Suppose one wishes to estimate the rate, *u*, of neutral mutations in a particular bacterial gene. We start by seeding a population with one bacterial cell and allowing exponential growth until there are *N* cells. We then measure the number of mutant cells, *m*, in the final population.

In this case, the existence of a mutation in a final cell is measured phenotypically by growing the cell under different conditions. For example, the initial environment may provide a particular nutrient, making a specific metabolic reaction within the cell unnecessary. When cells are subsequently grown in the absence of that nutrient, the previously unnecessary metabolic reaction becomes necessary for growth. Thus, we can infer mutations during the initial population growth that inactivate the metabolic function required during the subsequent phenotypic test.

The mutation rate is the number of mutational events divided by the number of cell divisions. If, for simplicity, we ignore cell death, then there are approximately *N* cell divisions to go from the initial cell to the final *N* cells.

Next, we need to estimate the number of mutational events. We measured *m* mutant cells among the final *N* cells. Different numbers of mutational events are consistent with a final number *m*. For example, a single mutational event that happened log(*m*) cell divisions back from the final cell would grow into *e*^log(*m*)^ = *m* final mutant cells. Or two mutational events that are log(*m*/2) cell divisions back from the final cells would also lead to *m* final mutant cells. Or there might be *m* mutational events, each occurring in the final cell division. Various combinations of mutational events and subsequent cellular growth would add to the same final value of *m*.

In other words, the number of final mutant cells provides some information about the number of mutational events but is not by itself sufficient to provide an exact value. We can gain more information by measuring the numbers of final mutant cells in several replicate populations.

For example, in Fig. 3, notice how the number of mutants at the median 0.5 level in height for each plot changes as *Nu* changes. If we estimate the median value for the number of mutants from a sample of populations, then we can then match that median to the value of *Nu* for a corresponding distribution. A median of 10^5^ would, for example, match *Nu* = 10000 in the right panel of Fig. 3. We know the value of *N*, so we can infer the value of *u*. Using the median gives a good rough approximation.

Other approaches may be more practical or provide more information. However, the idea is the same. In all cases, we are comparing observed measures for the number of mutants in the final population to the theoretical distribution based on the Luria–Delbrück process.^7,19^

## Genetics of human populations

Suppose we want to estimate how many mutations have occurred in a particular human population. For a focal stretch of DNA, we can measure how many of those sequences carry a mutation in the current population. Similarly, we can think of recombination events as changes that we count as mutants. For example, Hästbacka et al.^20^ estimated the amount of recombination and linkage in a DNA region that contains the diastrophic dysplasia (DTD) gene by studying the modern Finnish population.

In such applications to a single human population, we can obtain only a single measurement of the number of mutants or the number of recombinants. How much information does a single sample contain? In theory, how much more precise would estimates be if we could obtain additional independent samples? Evaluating these questions provides broad insight into what we may expect to see in natural populations and in how much information we can obtain during particular kinds of studies.

Hästbacka et al.^20^ used Luria–Delbrück theory to analyze the average number of events in their study population. However, as we discussed in prior sections, the average often poorly represents the Luria– Delbrück process because of the occasional very large values. Instead, analyzing the most likely value of *Nu* given the data provides a better approach. We typically know *N*, so an estimate of *Nu* provides an estimate of *u*, the mutation rate.

## Likelihood

Statistical tools for likelihood analysis of Luria– Delbrück processes have been given in the literature.^21,22^ However, the prior literature did not have an explicit form of the Luria–Delbrück distribution to work with. We can take advantage of the new Fréchet approximation in eqn 3. The amount of error from that approximation when used in likelihood analysis has not yet been quantified. For now, this presentation is best regarded as a conceptual introduction rather than an updated statistical procedure.

To develop likelihood analysis, we must rewrite the Fréchet cumulative density function in eqn 3 as a probability density function, which simply means that we need the derivative of *F*(*m*) with respect to *m*, written as

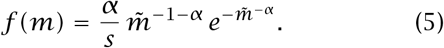

All of the parameters can be expressed in terms of the single parameter, *Nu*, as in eqn 4. Suppose we have data on the numbers of mutants in different samples, *m*_1_, *m*_2_,…. Then the log-likelihood of a particular parameter estimate, *Nu*, given the data, is

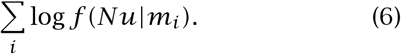

For example, if we have the single observation *m*_1_ = 10^5^ mutants, then Fig. 4a shows the log-likelihood for *Nu*. The relatively long left tail arises from the fact that smaller values of the mutation rate, *u*, occasionally give rise to large values of *m*, whereas bigger values of *u* rarely give rise to smaller values of *m*. Thus, the information in an observed value of *m* sets a strong upper bound on *u* but does not strongly bound lower values of *u*.

**Figure 4:**
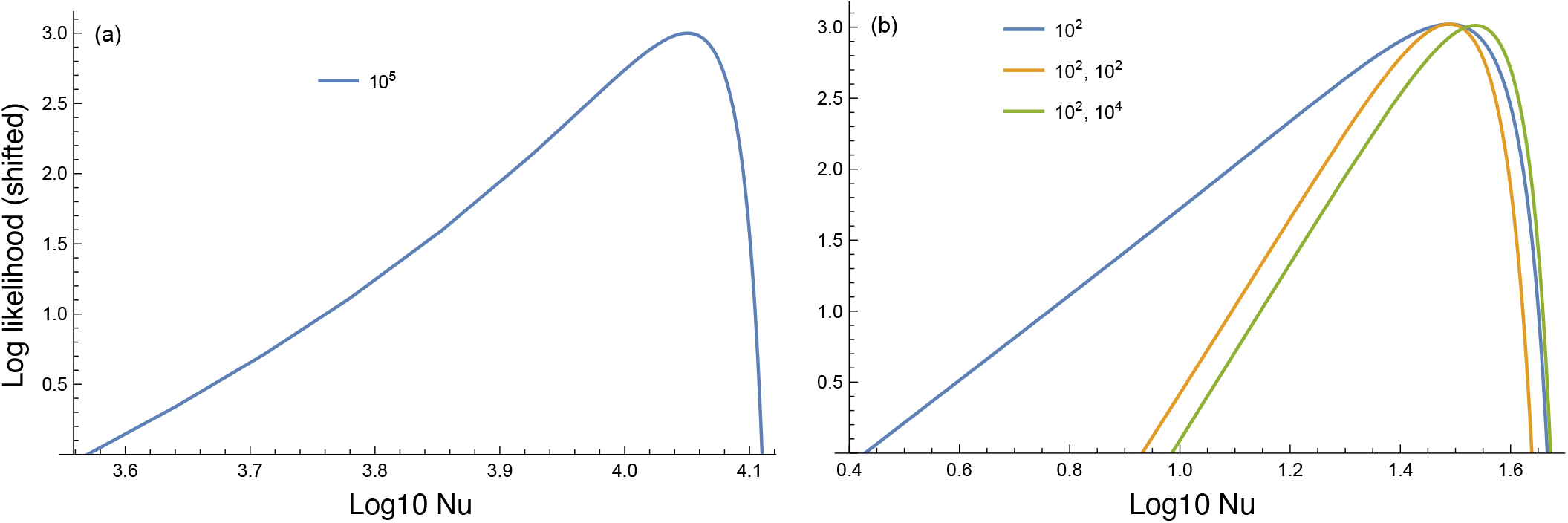
Log-likelihood for the parameter *Nu* given the values of the data shown in the legends of each plot. A constant was added to all log-likelihood values to shift the curves up so that the peaks are at a value of three. The zero values give the log-likelihood confidence intervals for which the most likely value is *e*^3^ ≈ 20 times more likely that the lowest values shown at the edges of the intervals. (a) For a single observation of *m* = 10^5^, the most likely value is approximately *Nu* = 10^4.05^. (b) Log-likelihood curves for different data combinations. The first (blue) curve has a single observed value, whereas the other two curves arise from pairs of observed values. These likelihood curves derive from the Fréchet approximation in eqn 3.

Figure 4b illustrates three interesting points. First, for the blue curve associated with the single observed value 10^2^, the range of *Nu* values is broader than for a single observed value 10^5^. Smaller numbers of mutational events cause greater variation, with most populations having few final mutants and a few populations having a lot of mutants.

Second, two observations of 10^2^ shrink the range of estimated values for a given confidence level by a bit less than one-half.

Third, comparing the pair of observations (10^2^,10^2^) versus (10^2^,10^4^), the estimated values change very little. The smaller observed value dominates the information. Once again, a small mutation rate can be highly consistent with a large observed value for the number of mutants, whereas a large mutation rate is weakly consistent with a small observed value for the number of mutants.

## Somatic mosaicism and disease

Animal bodies often arise from a single ancestral zygote. Cell division produces a large cellular population. The abstract Luria–Delbrück process provides a starting point for thinking about how mutations accumulate in such bodies, creating somatic mosaicism. Although actual development is more complex than the simple model, we can get a rough sense for numbers of mutant cells.

Prior articles have reviewed links between mosaicism and disease.^23–28^ Here, I focus on key conceptual issues.

### Mutant cells and the risk of disease

Cancer typically begins in a small piece of tissue. A local tumor develops and then disperses to cause widespread disease. In this case, each small piece of tissue that carries mutant cells poses an independent risk. The overall risk of cancer later in life rises roughly in proportion to the number of mutant cells produced during development through a Luria– Delbrück process.^29^

How does the somatic mosaicism induced by the Luria–Delbrück process affect other diseases? It depends on the mechanisms that link the origin of disease to the onset of widespread symptoms.^23,30^

Consider neurodegeneration. Certain inherited mutations predispose to disease.^31^ An inherited mutation is in every cell. What happens if the mutation is in 10% of cells, or 1% of cells, or 0.001% of cells? Can neurodegeneration start in a small local piece of tissue and then spread widely? If so, then a very small fraction of mutant cells could be associated with the origin the disease. The more mutant cells, the greater the chance that disease starts locally and spreads.

Ten or more years ago, just a few people emphasized this idea of local origin and subsequent spread for neurodegeneration.^32–34^ That idea for the origin of disease was then linked to somatic mosaicism.^23^ In recent years, the link between somatic mosaicism by a Luria–Delbrück process and neurodegeneration has motivated various studies and positive commentary.^27,28,35^ However, relative importance for that link remains an open problem.

Atherosclerosis provides another interesting example. That disease begins with aberrant physiological changes and tissue expansion in plaques that line the arteries.^36^ How much of atherosclerotic disease arises from Luria–Delbrück processes and somatic mosaicism? Current data provide clues but no clear answer.^37–39^

For other diseases, the questions are the same.^30^ To what extent could local changes in small pieces of tissue initiate widespread dissemination of disease? What role do mutations play in the risk of those initiating local tissue changes? Can local changes in hormone-secreting tissues cause disseminated disease?

In some cases, mutations may arise early in development, causing a high frequency of mutant cells. Such individuals may have disease phenotypes similar to cases of inherited disease. However, when tested genetically by analysis of a blood sample, some of those individuals will not have the mutations in their blood cells. They may be scored genetically as noninherited cases of disease but evaluated clinically with the symptoms of inherited cases.

### Variation between individuals

How much of the variation in disease risk between individuals arises from somatic mosaicism? In theory, the Luria–Delbrück process can cause significant variation in the somatic mutation burden for particular tissues when comparing between individuals. Empirically, it remains challenging to measure that variation and link the variation to disease. It may be particularly interesting to analyze those individuals who present with disease onset and symptoms typically associated with an inherited mutation but who lack the inherited mutation. The recent increase in interest for such problems and the advances in single cell genomics may eventually provide insight into these issues.

## Conclusions

This article provided a simple intuitive introduction to the Luria–Delbrück process and some of its consequences. The large literature includes many other applications and developments of theory to cover a variety of more realistic or complex assumptions.^7,40–45^

## Acknowledgments

The Donald Bren Foundation, National Science Foundation grant DEB-1939423, and DoD grant W911NF2010227 support my research.

